# Agreement between the ventilated capsule and the KuduSmart® device for measuring sweating responses to passive heat stress and exercise

**DOI:** 10.1101/2023.04.03.534973

**Authors:** Nicholas Ravanelli, Douglas Newhouse, Fergus Foster, Aaron R. Caldwell

**Affiliations:** School of Kinesiology, Lakehead University, Thunder Bay, Ontario, Canada; Centre for Research in Occupational Safety and Health, Laurentian University, Sudbury, Ontario Canada; Natick, MA

## Abstract

The present study assessed agreement between the KuduSmart® device and the ventilated capsule (VC) technique for measuring: i) minute-averaged local sweat rate (LSR); ii) sweating onset; iii) thermosensitivity; and iv) steady-state LSR, during passive heat stress and exercise. On two separate occasions for each heat stress intervention, participants were either passively heated by recirculating hot water (49°C) through a tube-lined garment until rectal temperature increased 1°C over baseline (n=8), or walked on a treadmill for 60 minutes at a fixed rate of heat production (n=9). LSR of the forearm was concurrently measured with a VC and the KuduSmart® device secured within ∼2 cm. Mean body temperature was calculated as the weighted average between rectal (80%) and mean skin (20%) temperature. Using a ratio scale Bland-Altman analysis with the VC as the reference, the KuduSmart® device demonstrated systematic bias for minute-averaged LSR (1.19 [1.09, 1.28]) and steady-state LSR (1.16 [1.09,1.23]), however no bias for thermosensitivity (1.08 [1.00, 1.16]) or sweating onset (1.00 [1.00, 1.00]). Although poor agreement was observed for minute-averaged LSR (CV = 43.3%), more than 73% of all observations with the KuduSmart® device (n=2795) were within an absolute error of < 0.2 mg/cm^2^/min to the VC. The KuduSmart® device demonstrated acceptable agreement for steady-state LSR (CV = 19.5%) and thermosensitivity (CV = 21.7%), and almost perfect agreement for sweating onset (CV = 0.14%). Collectively, the KuduSmart® device may be a satisfactory in-field solution for assessing the sweating response to heat stress.

## Introduction

The secretion of sweat onto the skin surface is a necessary heat loss mechanism to mitigate deleterious rises in core temperature during heat stress. Researchers interested in thermoregulatory responses during heat stress typically measure local sweat rate to assess sudomotor output. One of the most common lab-based methods is the ventilated capsule technique which estimates local sweat rate by measuring the difference in water vapour content of air entering and exiting a capsule affixed to the skin. While this technique has been employed for over 60 years (Bullard 1962), no commercially available ventilated capsule system exists requiring researchers to curate custom solutions using hygrometers and flow meters interfaced with a data acquisition unit. Further, these custom solutions often require pressurized gas cylinders with a known water vapor content to supply the effluent air. While this setup is ideal for lab-based settings, it is impractical for field use.

The technical absorbent patch is one of the more common methods for measuring local sweat rate in the field (Havenith et al. 2008). In brief, a highly absorbent patch is placed on the skin surface for a fixed amount of time providing an index of local sweat rate; however, this method lacks the temporal resolution offered from a ventilated capsule. Alternatively, patches have been developed that allow end-users to measure the speed of sweat accumulation within microfluidic channels (Nyein et al. 2021; Ghaffari et al. 2023), or assess changes in conductance as channels gradually fill (Steijlen et al. 2022). However, the volume capacity of the channel limits its application for extended use or at higher sweat rates. More recently, a promising new portable capsule design was proposed where air is continuously circulated through silica gels to absorb moisture and the microenvironment of the skin surface is measured using a thermohygrometer positioned in proximity to the circulating microfan inlet (Uchida et al. 2022). Unfortunately, this closed-loop system will be restricted by the maximum water-carrying capacity of the silica gels within the upper compartment which limits its applicability for measuring high sweat rates for prolonged periods of time. Moreover, as described by the authors, the aforementioned portable capsule was tethered to two additional modules for offline data logging, with no details describing if and how the device was externally powered.

In contrast to previously mentioned in-field techniques for measuring local sweat rate, the KuduSmart® device (developed by Crossbridge Scientific Ltd. UK) employs an open loop system similar to the traditional ventilated capsule technique whereby air is drawn through a single channel on the skin surface using a microfan. In comparison to the closed loop system by Uchida et al. (2022), both influent and effluent air are sampled independently by a thermohygrometer and the net difference in water vapour provides an index of local sweat rate. Previous work by Relf et al. (2019) observed a moderate interclass correlation coefficient (0.88) for test-retest reliability of the average local sweat rate over 30-minutes of cycling using the KuduSmart® device. In a separate study, Relf et al. (2020) concluded that the KuduSmart® device detected an increased average local sweat rate during 30-minutes of cycling at 3.5 W/kg in 35°C and 50%RH following heat acclimation, likely due to the earlier onset of sweating and higher sweat rates (Ravanelli et al. 2019). In addition to monitoring real-time data collection on a Bluetooth connected tablet or smartphone, the KuduSmart® device has a relatively high temporal resolution (0.17Hz, or samples every 6 seconds) positioning it as a possible substitution to the traditional ventilated capsule technique for field- or lab-based studies. The purpose of the present study was to evaluate the agreement between the KuduSmart® device and the ventilated capsule technique for measuring minute-averaged local sweat rate and assessing key sweating characteristics such as sweat onset, thermosensitivity, and steady-state local sweat rate during passive heat stress and exercise.

## Methods

### Ethical approval

Ethics approval was received by the Lakehead University Research Ethics Board and the Thunder Bay Regional Health Sciences Centre Research Ethics Board for the passive heat stress and exercise arms of the study, respectively. The data presented in this study are from secondary analysis of larger independent studies satisfying unique objectives. The study conformed to the standards set by the Declaration of Helsinki, except for database registration. All participants provided verbal and written consent prior to participating.

### Participants

A total of 15 participants completed either the passive heat stress or exercise arm of the present study (6 females, 25.0±4.7 y, 1.8±0.1 m, 74.8±9.3 kg), with 2 participants completing both arms (1 male and 1 female). As the primary aim of the current study was to evaluate the agreement between two measurement techniques, the menstrual cycle for all female participants was not standardized.

### Study Design

Prior to all visits, participants were instructed to drink ∼500 mL water and eat a light snack before arrival, and avoid coffee, alcohol, and strenuous exercise for 12 hours. For either passive heat stress or exercise, time of day for each trial was matched within participant.

### Passive heat stress protocol

8 participants completed 2 passive heat stress sessions. Upon arrival to the laboratory, participants were instrumented and donned a water-perfused suit (COOLTUBEsuit, Med-Eng, LLC). Following instrumentation, participants rested in a supine position for 10 minutes while 32°C water (TC-102D-115, Brookfield) was circulated through the suit. Afterwards, the water perfusing the suit was increased to 49°C and maintained until rectal temperature increased by 1°C above baseline.

### Exercise protocol

9 participants completed 2 exercise sessions. Upon arrival to the laboratory, participants were instrumented and rested in a seated position for ∼20 minutes in a climate-controlled room (∼29C, 30%RH). Following the rest period, participants began a 60-minute treadmill march at a fixed rate of heat production (∼550W).

### Instrumentation

Mean skin temperature was calculated using a 4-point ((Ramanathan 1964) passive heat stress) or 14-point ((Parsons 2014), exercise) weighted average sampled at 0.2Hz using Thermocron iButtons (DS1922L-F5, Maxim Integrated) affixed to the skin using surgical tape. Rectal temperature was measured using a general-purpose thermistor probe (9fr, DeRoyal) self-inserted in private 12 cm past the anal sphincter and sampled in real-time using a data acquisition unit at 100 Hz (PowerLab 8/35, AD Instruments). Mean body temperature was calculated using a weighted average of rectal (80%) and mean skin (20%) temperature and averaged every minute. A ventilated capsule (surface area coverage = 2.89cm^2^) was placed on anterior aspect of the left forearm and supplied with influent dry gas governed at 1.7 L/min using a glass flowmeter (FL3905G, Omega Engineering). Effluent air from the ventilated capsule was measured using a temperature and humidity probe (HMT330, Vaisala) with the analog outputs connected directly to the data acquisition unit. Local sweat rate within the ventilated capsule was calculated as the product of air flow and the net difference in absolute water vapour content between effluent and influent air, normalized to capsule surface area and expressed in mg/cm^2^/min. The KuduSmart® device (Crossbridge Scientific Ltd., UK) was placed within ∼2 cm of the ventilated capsule (See Figure 1), and tightness of the strap was replicated within person. Placement of the KuduSmart® device on the forearm was counterbalanced between trials within person for proximal/distal placement relative to the ventilated capsule. Raw Kudu scores (an arbitrary unit quantifying the relative humidification of the effluent air) were broadcasted every 6 seconds via Bluetooth to the KuduSmart® Android application on the supplied tablet (Lenovo). The Kudu score was multiplied by a conversion factor provided by the manufacturer (0.26) to produce local sweat rate values in mg / cm^2^ /min. For each measurement technique, the onset of sweating and sudomotor thermosensitivity was determined objectively using segmented linear regression analysis (GraphPad Prism 9.4, la Jolla, USA) of minute-averages of local sweat rate against mean body temperature.

**Figure 1.**
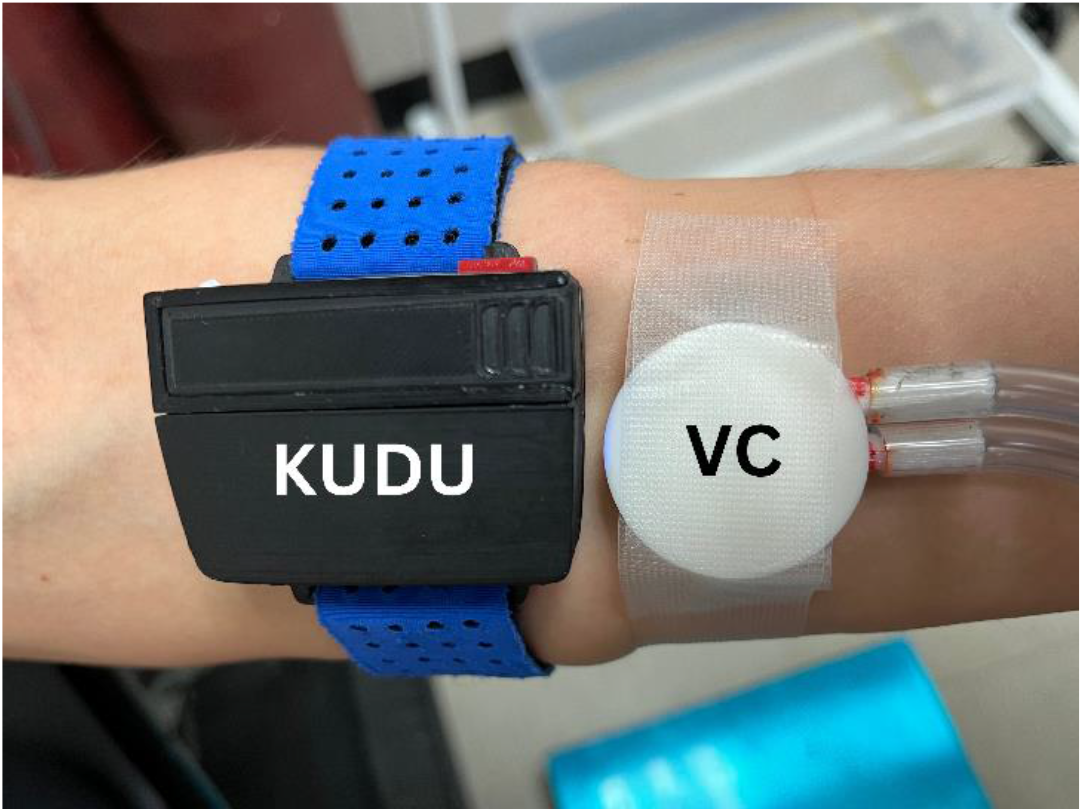
Placement of the KuduSmart® (KUDU) device and the ventilated capsule (VC) to concurrently measure forearm local sweat rate.

### Statistical Analysis

As the primary purpose of the present study was to determine the agreement between the KuduSmart® device and the ventilated sweat capsule technique, all data from passive heat stress and exercise tests were combined for analysis. The following characteristics of sweating were assessed for agreement between both techniques: i) minute-averaged local sweat rate, ii) the mean body temperature corresponding to the onset of sweating, iii) the thermosensitivity of local sweat rate to changes in mean body temperature, and iv) the average local sweat rate during the final 15 minutes where a visible steady-state had been achieved. Limits of agreement were calculated using the SimplyAgree package (Caldwell 2022) which accounts for the nested nature of the data (Zou 2013). Due to measurement error appearing to be proportional to the mean (Bland and Altman 1996), all measures were log-transformed and the limits of agreement were derived from a ratio scale. The coefficient of variation was computed for each of the sweating metrics evaluated.

## Results

Mean bias and limits of agreement from the Bland-Altman analysis and coefficient of variation are presented in Table 1, and the data is visualized against a line-of-identity in Figure 2. For minute-averaged local sweat rate, the Bland-Altman analysis suggested proportional bias and the coefficient of variation highlighting poor agreement between the KuduSmart® device and the ventilated capsule technique. 73.63% of all observations (n=2795) from the KuduSmart® device were no more than 0.2 mg/cm^2^/min different to the time-matched minute-averaged local sweat rate measured from the ventilated capsule (Figure 3). The KuduSmart® device demonstrated good agreement with no proportional bias relative to the ventilated capsule for detecting the onset of sweating. While no proportional bias was observed, the KuduSmart® device demonstrated acceptable agreement for assessing thermosensitivity. Lastly, acceptable agreement was observed between the KuduSmart® device and the ventilate capsule technique for measuring steady-state sweating, however proportional bias was observed.

**Table 1.**
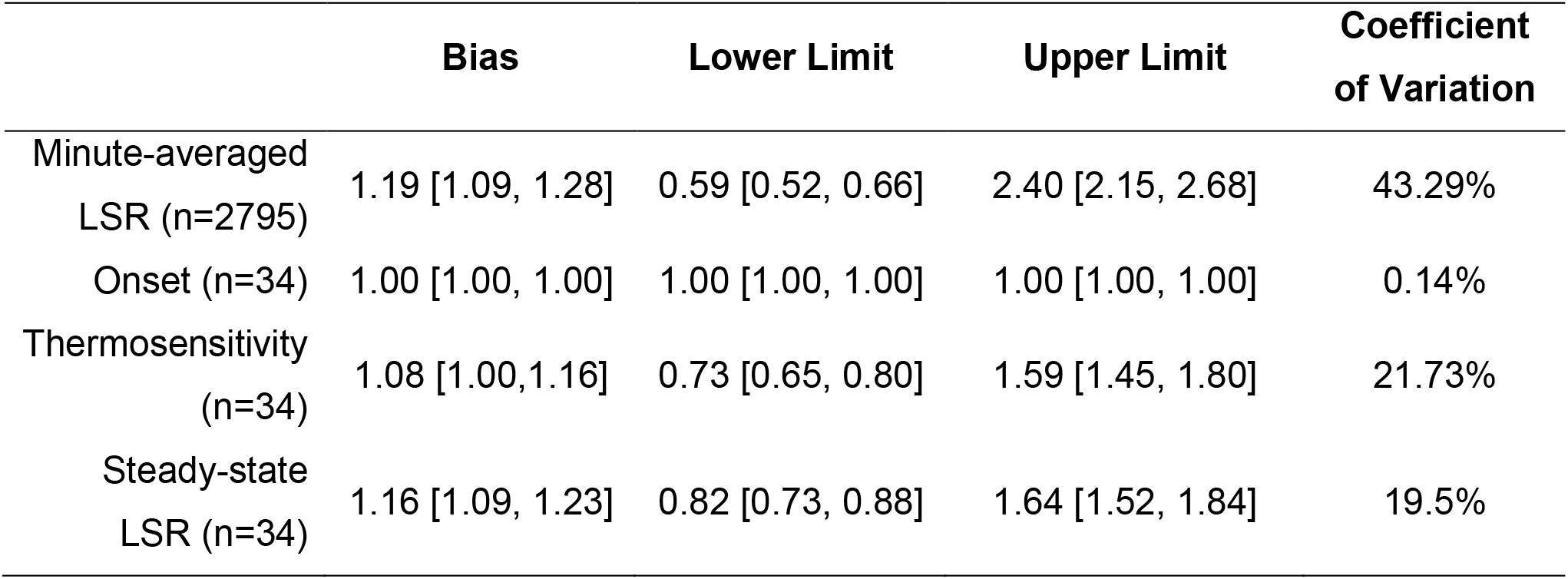
Mean bias and limits of agreement with 95% confidence intervals from the Bland-Altman analysis for assessing the agreement between the KuduSmart® device and the ventilated capsule technique when measuring minute-averaged local sweat rate (LSR), sweating onset, thermosensitivity, and steady-state LSR. The number of individual observations for each parameter are presented as (n=observations).

|  | **Bias** | **Lower Limit** | **Upper Limit** | **Coefficient of Variation** |
| --- | --- | --- | --- | --- |
| Minute-averaged LSR (n=2795) | 1.19 [1.09, 1.28] | 0.59 [0.52, 0.66] | 2.40 [2.15, 2.68] | 43.29% |
| Onset (n=34) | 1.00 [1.00, 1.00] | 1.00 [1.00, 1.00] | 1.00 [1.00, 1.00] | 0.14% |
| Thermosensitivity (n=34) | 1.08 [1.00, 1.16] | 0.73 [0.65, 0.80] | 1.59 [1.45, 1.80] | 21.73% |
| Steady-state LSR (n=34) | 1.16 [1.09, 1.23] | 0.82 [0.73, 0.88] | 1.64 [1.52, 1.84] | 19.5% |

**Figure 2.**
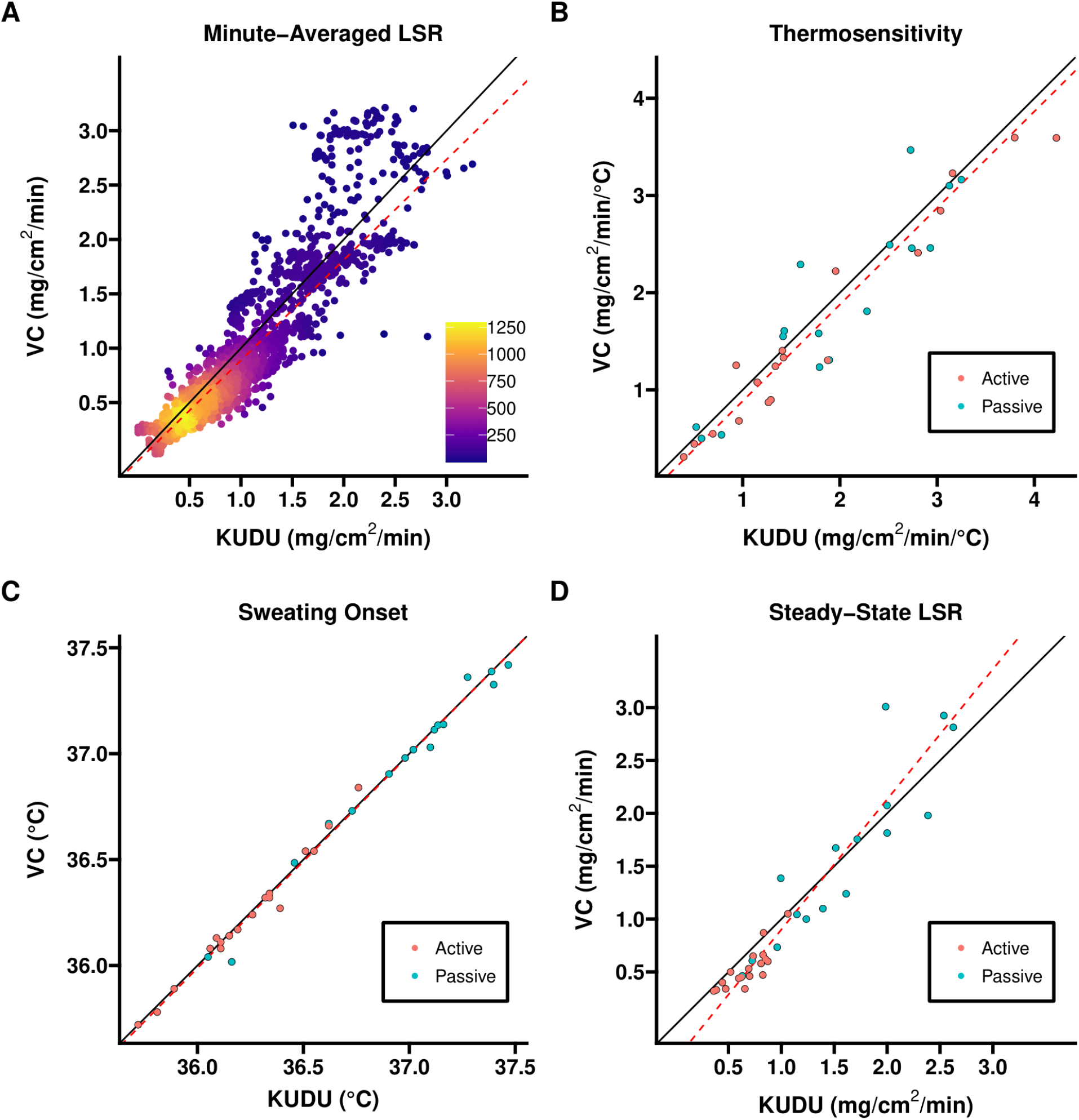
Line of identity plots (Solid black line) between the ventilated capsule (VC) and KuduSmart® device for measuring minute averaged local sweat rate (LSR, A), sudomotor thermosensitivity (B), the onset of sweating (C), and iv) steady-state LSR (D). The dashed red line denotes the results of a Weighted Deming Regression (See data repository; Caldwell and Ravanelli 2023).

**Figure 3.**
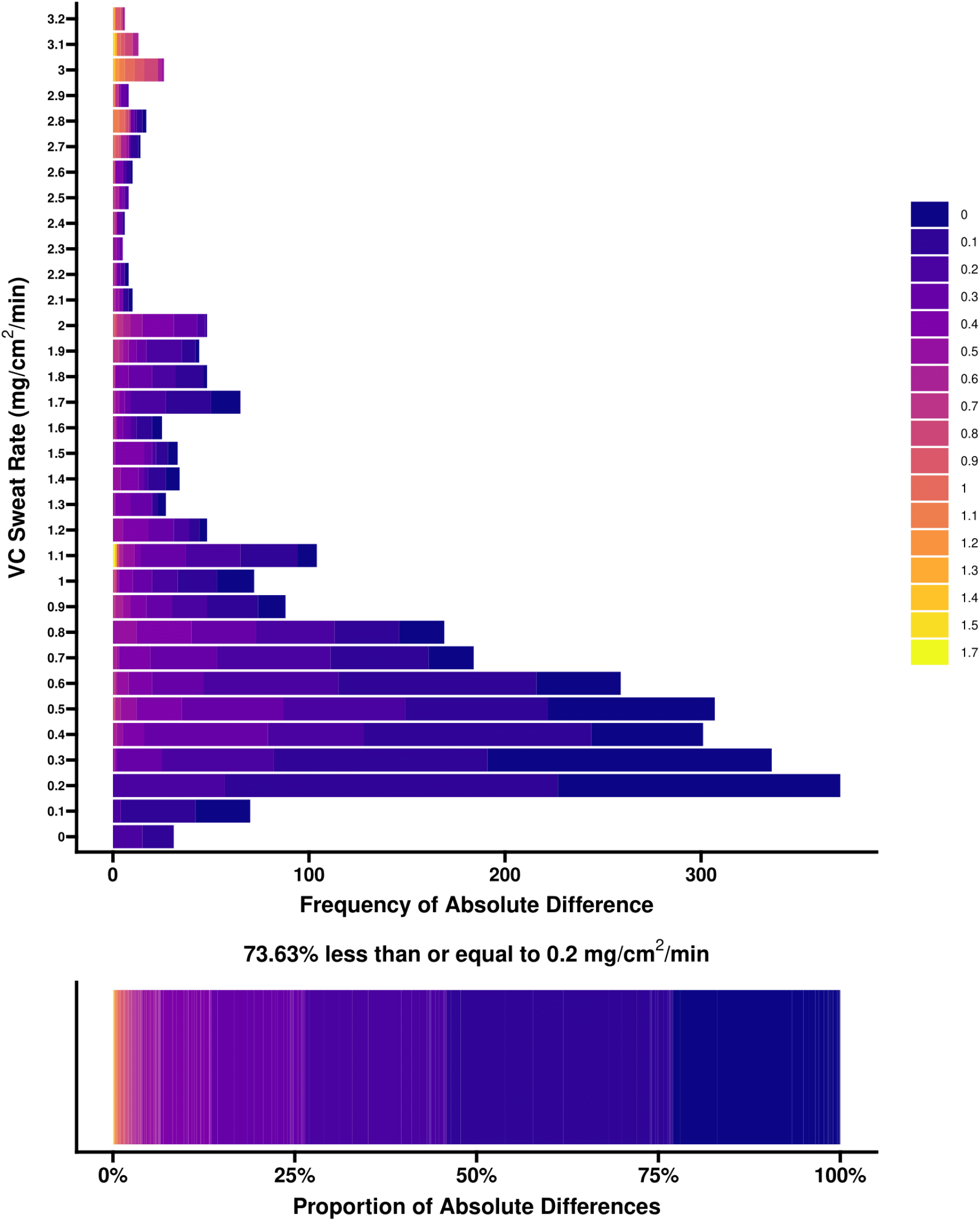
Frequency (top) and proportion (bottom) of absolute differences between the observed local sweat rate measured with the ventilated capsule (VC) and the KuduSmart® device (heat map).

## Discussion

The present study evaluated the agreement between the wireless KuduSmart® device and the VC technique during passive heat stress and exercise. The KuduSmart® device demonstrated near perfect agreement for detecting the onset of sweating with no proportional bias, acceptable agreement for assessing sweating thermosensitivity with no proportional bias, and acceptable agreement for steady-state local sweat rate with slight proportional bias. Despite the poor agreement and proportional bias for minute-averaged local sweat rate measurements, the absolute error of the KuduSmart® device relative to the VC was less than 0.2 mg/cm^2^/min for over 70% of the observations (Figure 3). Collectively, our findings provide support for the use of the KuduSmart® device for evaluating key characteristics of the sweating response during heat stress in conditions where the VC technique may be impractical or inaccessible.

Relf et al. (2019, 2020) provided initial evidence to support the validity and reliability of the KuduSmart® device for assessing average sweat rate over 30 minutes in comparison to the technical absorbent technique. We observed a higher coefficient of variation (19.5% vs 13.5%) with the KuduSmart® device in comparison to the VC technique when averaging the final 15 minutes of heat stress where a plateau in sweat rate was observed, across a wider range of local sweat rates (0.32 – 3.01 mg/cm^2^/min, Figure 2,3). As demonstrated in Figure 3, the absolute error of the KuduSmart® device increases at higher sweat rates which may explain the increased coefficient of variation observed in the present study, warranting further work. Nevertheless, the current study extends previous findings by demonstrating the KuduSmart® device’s capacity to detect the onset of sweating and assess thermosensitivity with acceptable agreement to the VC, key characteristics of the sudomotor response to heat stress that can be altered by various factors including, but not limited to, acclimation (Ravanelli et al. 2019; Barry et al. 2020), age (Stapleton et al. 2015), disease (Chaseling et al. 2021), and drugs (Crandall et al. 2002).

Advancements in wearable biosensors and wireless devices have enabled researchers to evaluate physiological responses in more ecologically valid conditions with similar measurement accuracy to lab-based equipment. More recent prototypes of a portable local sweat rate monitoring device have assumed influent air in a closed loop system (Uchida et al. 2022), which over time may become saturated and systematically underestimate the local sweat rate measured. While other devices and techniques have been explored to measure local sweat rate in the field (Havenith et al. 2008; Nyein et al. 2021; Steijlen et al. 2022; Ghaffari et al. 2023), to our knowledge, the KuduSmart® is the only device that operates similarly to the ventilated capsule technique without the need for an anhydrous gas supply, and communicates wirelessly over Bluetooth® to a companion application for real-time monitoring.

Based on the present findings, the KuduSmart® device could serve as a portable ‘*drop in*’ replacement to the traditional ventilated capsule technique offering researchers greater accessibility and flexibility in measuring sudomotor control during passive heat stress or exercise.

For example, this device could enable real-time monitoring of local sweat rate responses during occupational work in thermally stressful environments, during athletic training or competition, or over prolonged periods of exposure to extreme heat. Further, the KuduSmart® device may help clinicians and researchers characterize how various factors influence sudomotor output including, but not limited to, aging, comorbidity, medications, fitness, and heat acclimation. The replaceable battery that is easily accessible while being worn offers the potential for prolonged use in excess of the estimated ∼90-minute runtime from a single AAA battery. However, it remains unclear whether transient changes in the ambient conditions, a more humid environment, or clothing overtop, may independently affect the accuracy of the KuduSmart® device in measuring local sweat rate and evaluating sweating response to heat stress.

## Conclusion

In comparison to the ventilated capsule technique, the portable KuduSmart® device demonstrated near perfect agreement for measuring sweating onset, acceptable agreement for thermosensitivity and steady-state local sweat rate, and poor agreement for minute-averaged local sweat rate. Collectively, the KuduSmart® device enables researchers to evaluate key characteristics of the sweating response to heat stress where a ventilated capsule setup may be impractical.

## Author contribution statement

NR was responsible for conceptualization, data curation, funding acquisition, methodology, project administration, validation, supervision, writing original draft, and writing – review and editing. DN & FF were responsible for conceptualization, methodology, project administration, and formal analysis, writing – review and editing. AC was responsible for formal analysis, data curation, methodology, visualization, and writing – review and editing.

## Competing interests statement

The authors declare there are no competing interests.

## Funding

This research was support by Dr. Nicholas Ravanelli’s NSERC Discovery Grant (grant no. RGPIN-2022-05096), and a Centre for Research in Occupational Safety and Health Seed Grant.

## Data Availability statement

Data generated or analyzed during this study are available in the Open Source Framework repository (DOI: 10.17605/OSF.IO/7Y8MR; Caldwell and Ravanelli 2023).

## Notes

### Competing Interest Statement

The authors have declared no competing interest.

## References

Barry, H., Chaseling, G.K., Moreault, S., Sauvageau, C., Behzadi, P., Gravel, H., Ravanelli, N., and Gagnon, D. 2020. Improved neural control of body temperature following heat acclimation in humans. The Journal of Physiology 598(6): 1223–1234. doi:10.1113/JP279266.

Bland, J.M., and Altman, D.G. 1996. Measurement error proportional to the mean. BMJ 313(7049): 106–106. doi:10.1136/bmj.313.7049.106.

Bullard, R.W. 1962. Continuous recording of sweating rate by resistance hygrometry. Journal of Applied Physiology 17(4): 735–737. doi:10.1152/jappl.1962.17.4.735.

Caldwell, A.R. 2022. SimplyAgree: An r package and Jamovi module for simplifying agreement and reliability analyses. Journal of Open Source Software 7(71): 4148. doi:10.21105/joss.04148.

Caldwell, A.R., and Ravanelli, N. 2023. Data and code for Agreement between the ventilated capsule and the KuduSmart® device for measuring sweating responses to passive heat stress and exercise. OSF. doi:10.17605/OSF.IO/7Y8MR.

Chaseling, G.K., Filingeri, D., Allen, D., Barnett, M., Vucic, S., Davis, S.L., and Jay, O. 2021. Blunted sweating does not alter the rise in core temperature in people with multiple sclerosis exercising in the heat. American Journal of Physiology-Regulatory, Integrative and Comparative Physiology 320(3): R258–R267. American Physiological Society. doi:10.1152/ajpregu.00090.2020.

Crandall, C.G., Vongpatanasin, W., and Victor, R.G. 2002. Mechanism of cocaine-induced hyperthermia in humans. Ann Intern Med 136(11): 785–791. American College of Physicians. doi:10.7326/0003-4819-136-11-200206040-00006.

Ghaffari, R., Aranyosi, A.J., Lee, S.P., Model, J.B., and Baker, L.B. 2023. The Gx Sweat Patch for personalized hydration management. Nat Rev Bioeng 1(1): 5–7. Nature Publishing Group. doi:10.1038/s44222-022-00005-5.

Havenith, G., Fogarty, A., Bartlett, R., Smith, C.J., and Ventenat, V. 2008. Male and female upper body sweat distribution during running measured with technical absorbents. Eur J Appl Physiol 104(2): 245–255. doi:10.1007/s00421-007-0636-z.

Nyein, H.Y.Y., Bariya, M., Tran, B., Ahn, C.H., Brown, B.J., Ji, W., Davis, N., and Javey, A.2021. A wearable patch for continuous analysis of thermoregulatory sweat at rest. Nat Commun 12(1): 1823. Nature Publishing Group. doi:10.1038/s41467-021-22109-z.

Parsons, K. 2014. Human Thermal Environments : The Effects of Hot, Moderate, and Cold Environments on Human Health, Comfort, and Performance, Third Edition. CRC Press. doi:10.1201/b16750.

Ramanathan, N.L. 1964. A new weighting system for mean surface temperature of the human body. J Appl Physiol 19(3): 531–533.

Ravanelli, N., Coombs, G., Imbeault, P., and Jay, O. 2019. Thermoregulatory adaptations with progressive heat acclimation are predominantly evident in uncompensable, but not compensable, conditions. Journal of Applied Physiology 127(4): 1095–1106. doi:10.1152/japplphysiol.00220.2019.

Relf, R., Eichhorn, G., Waldock, K., Flint, M.S., Beale, L., and Maxwell, N. 2020. Validity of a wearable sweat rate monitor and routine sweat analysis techniques using heat acclimation. Journal of Thermal Biology >90: 102577. doi:10.1016/j.jtherbio.2020.102577.

Relf, R., Willmott, A., Flint, M.S., Beale, L., and Maxwell, N. 2019. Reliability of a wearable sweat rate monitor and routine sweat analysis techniques under heat stress in females. Journal of Thermal Biology 79: 209–217. doi:10.1016/j.jtherbio.2018.12.019.

Stapleton, J.M., Poirier, M.P., Flouris, A.D., Boulay, P., Sigal, R.J., Malcolm, J., and Kenny, G.P. 2015. At what level of heat load are age-related impairments in the ability to dissipate heat evident in females? PLoS ONE 10(3): e0119079. doi:10.1371/journal.pone.0119079.

Steijlen, A.S.M., Jansen, K.M.B., Bastemeijer, J., French, P.J., and Bossche, A. 2022. Low-cost wearable fluidic sweat collection patch for continuous analyte monitoring and offline analysis. Anal. Chem. 94(18): 6893–6901. American Chemical Society. doi:10.1021/acs.analchem.2c01052.

Uchida, K., Ogawa, Y., Kataoka, Y., Manabe, K., Aida, T., Kamijo, Y., Takahashi, S., Ikefuchi, R., Nose, H., and Masuki, S. 2022. New portable device for continuous measurement of sweat rate under heat stress during field tests. Journal of Applied Physiology 132(4): 974–983. American Physiological Society. doi:10.1152/japplphysiol.00155.2021.

Zou, G. 2013. Confidence interval estimation for the Bland–Altman limits of agreement with multiple observations per individual. Stat Methods Med Res 22(6): 630–642. doi:10.1177/0962280211402548.

